# Vertical transmission of *Borrelia turicatae* (Spirochaetales: *Borreliaceae*) by autogenously reproducing *Ornithodoros turicata* (Ixodida: Argasidae) female naturally infected with the spirochetes

**DOI:** 10.1101/2023.06.26.546580

**Authors:** Serhii Filatov, Aparna Krishnavajhala, Job E. Lopez

**Affiliations:** Department of Pediatrics, National School of Tropical Medicine, Baylor College of Medicine, Houston, TX, USA; Department of Molecular Virology and Microbiology, Baylor College of Medicine, Houston, TX, USA

**Author notes:** Address correspondence to Job E. Lopez. Baylor College of Medicine, One Baylor Plaza, Houston, TX 77030.

## Abstract

*Ornithodoros turicata* is a vector of relapsing fever (RF) spirochetes in North America and transmits *Borrelia turicatae* to a variety of vertebrate hosts. The remarkably long lifespan of *O. turicata* and its ability to maintain spirochetes horizontally (between life stages) and vertically to progeny promotes the perpetuation of *B. turicatae* in nature. Nevertheless, the reproductive biology of *O. turicata* is poorly understood. In this report, we collected ticks from a park within a neighborhood of Austin, Texas. They were reared to adulthood and male ticks were individually housed with females. We observed autogenous reproduction by the ticks and further investigated vertical transmission of *B. turicatae* by quantifying filial infection rates in a cohort of progeny ticks. These results indicate that *O. turicata* transovarially transmits *B. turicatae* during autogenous reproduction and further signify the tick as a natural reservoir of the spirochetes.

**Importance:** Previous research has implicated *Ornithodoros* ticks, including *Ornithodoros turicata*, as long-term reservoirs of relapsing fever (RF) spirochetes. Considering the tick’s long lifespan and their efficiency in maintaining and transferring spirochetes within the population, the infection could persist in a given enzootic focus for decades. However, little is known about the relative importance of horizontal and vertical transmission routes in the persistence and evolution of RF *Borrelia*. Our observations on the reproductive biology of *O. turicata* in the absence of vertebrate hosts indicate an additional mechanism by which *B. turicata* can be maintained in the environment. This work establishes the foundation for studying *O. turicata* reproduction and spirochete-vector interactions, which will aid in devising control measures for *Ornithodoros* ticks and RF spirochetes.

## Introduction

The soft tick *Ornithodoros turicata* is a major vector of relapsing fever (RF) spirochetes in North America and is commonly found throughout the southern United States into Mexico (1). Wherever the vector occurs, enzootic foci can be formed between the pathogenic spirochete, *Borrelia turicatae*, and their wildlife hosts. Moreover, spillover events occur when humans or companion animals enter habitats infested with infected ticks. For example, recent work indicates that *B. turicatae* is an emerging threat in parks and recreational areas in highly populated cities of Texas (2-4). To further understand the public health impact of *B. turicatae*, knowledge of the tick’s biology and pathogen-vector interactions is needed.

The life cycle of *O. turicata* is complex and the dynamics of vertical transmission of *B. turicatae* are poorly understood. The life span of *O. turicata* nears 10 years (5), and adult female ticks can oviposit multiple times (6, 7). Gordon Davis reported a cohort of field collected *O. turicata* females oviposited five times in a 12-month period (7). Once eggs hatch, *O. turicata* has upwards of five nymphal instar stages (5). After molting into adults, vertically infected ticks can maintain *B. turicatae* for at least five generations (7). Collectively, prior work indicates that infected *O. turicata* can bypass the need to acquire the infection from a vertebrate host and serves as a reservoir for *B. turicatae*.

An interesting nuance of *Ornithodoros* reproduction that should be considered in combination with transovarial transmission is autogeny. This is the ability to produce eggs without the need for a blood meal. While known to occur in *Ornithodoros* ticks, autogeny varies between species (8). Endris and colleagues reported that during two years of rearing *Ornithodoros puertoricensis* they failed to observe autogeny (9). Alternatively, autogeny has been reported in species like *Ornithodoros tholozani* and *Ornithodoros fonsecai* (8, 10). However, we have not found a report characterizing autogeny in *O. turicata*.

The current study was based on a serendipitous observation while assessing *O. turicata* reproduction in a group of naturally infected ticks. Interestingly, as the nymphs molted into adults, we observed cases of autogeny. In one cohort of ticks, we determined rates of transovarial transmission to the F1 progeny. Our work indicates that blood feeding in the first gonotrophic cycle is not essential for *O. turicata* female reproduction and that transovarial transmission occurs by autogeny.

## Results

### Collection of ticks from a park in Austin, Texas

Tick collections were part of routine surveillance of *Ornithodoros* species in populated cities of Texas. A park was identified in a neighborhood of south Austin (Figure 1A). This ecoregion consisted of limestone outcroppings and Live Oak Mesquite Savanna (Figure 1B). As ticks were lured from outcroppings with dry ice as a source of carbon dioxide, they were collected and housed together (Figure 1C). We collected 41 ticks. Of these, 15 were adults and 26 were nymphs. Five nymphs died between the time the ticks were collected and evaluated in our laboratory.

**Figure 1.**
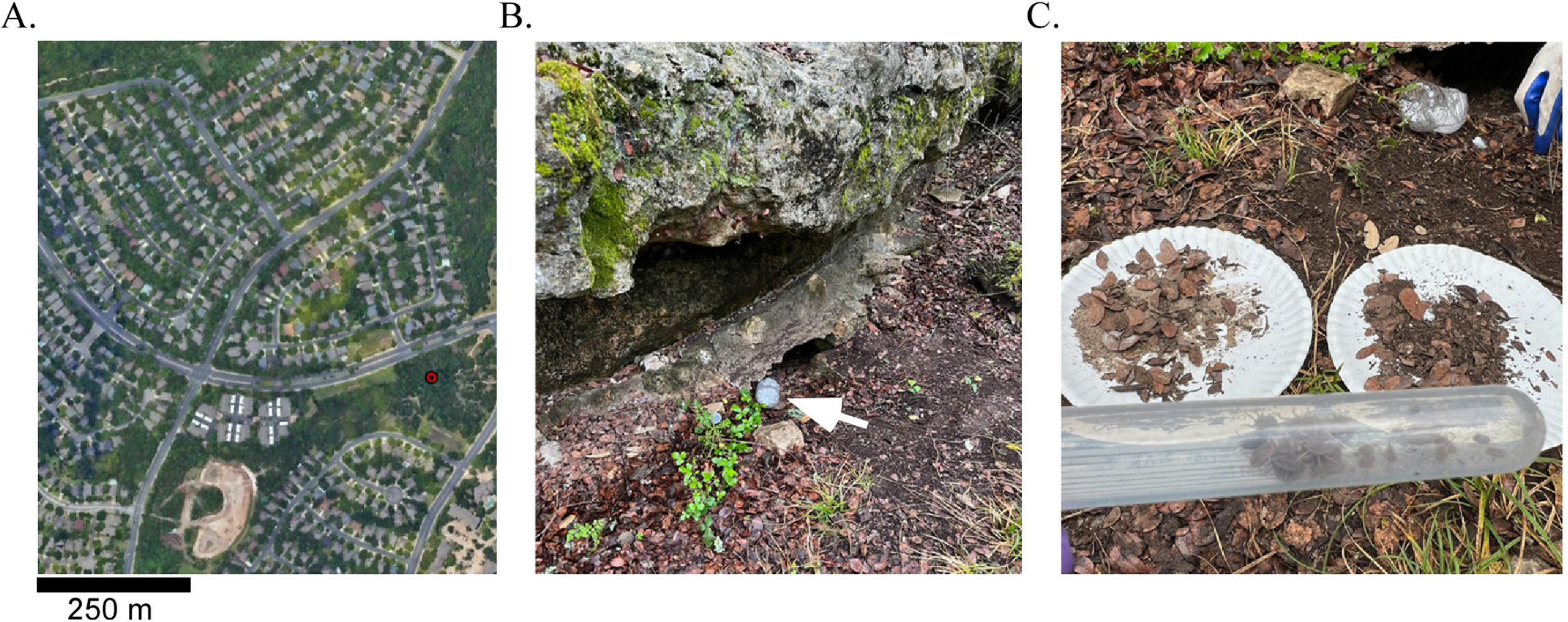
Collection site of *Ornithodoros turicata* in Austin, TX, USA. Shown is an aerial view (Google Earth©) of the collection site with a scale in the bottom left corner (A). The red dot represents the location where ticks were collected. A container with dry ice was placed by a limestone outcropping and used to bait ticks (B). The white arrow points to the dry ice container. Also shown is a 15 ml tube containing the ticks that were collected (C). Shown on the paper plates is the material from which ticks were harvested (C).

### Speciation of field collected ticks

In the laboratory we determined the species of field collected ticks by microscopy and through a molecular approach. Morphological characterization indicated that the ticks were *O. turicata*. For molecular typing, we amplified a portion of the *cox1* gene from two ticks, which produced an ∼700 nucleotide fragment, as determined by agarose gel electrophoreses. Sequencing and BLAST analysis determined 99.71% nucleotide identity with *O. turicata*. The sequences were deposited to GenBank under accession numbers OR047917 and OR047918. These findings indicated an additional endemic focus of *O. turicata* in Austin, Texas.

### Autogenous reproduction by *O. turicata*

Of the 21 live field-collected late-stage nymphs assessed, three died after blood feeding on laboratory mice and nine molted into females and nine into males. Interestingly, within 90 days after pairing males and females there were numerous larvae in four out of eight tubes. These findings indicated that *O. turicatae* females reproduced and laid eggs autogenously. We subsequently determined whether transovarial transmission occurred in one of the cohorts.

### Assessment of transovarial transmission

We reared larvae to the second instar stage and xenodiagnosis was performed to determine whether these ticks were infected with *B. turicatae*. Feeding F1 second instar stage nymphs on mice indicated that the ticks were infected (Table 1). We visualized spirochetes in the blood of five of seven animals by the fifth day after feeding ticks. Spirochetes were observed in the blood of another animal on the ninth day. The mouse in which we failed to detect spirochetes by microscopy seroconverted to *B. turicatae* protein lysates as determined by immunoblotting (Table 1). This indicated that the infection was below the limit of detection by microscopy.

**Table 1.** Assessment of murine infection and *B. turicatae* densities in autogenously produced ticks.

| Mouse # | Assessment of infection | | # of ticks infected/tested<sup>‡</sup> | $\bar{X}$ (SD) *flaB* copies per infected tick<sup>‡</sup> |
| --- | --- | --- | --- | --- |
|  | Microscopy (DPF) | Seroconversion by immunoblot |  |  |
| M1 | + (5) | ND | 3/15 | 2.28x10<sup>5</sup> (±8.2x10<sup>4</sup>) |
| M2 | - | + | 7/15 | 2.29x10<sup>5</sup> (±9.49x10<sup>4</sup>) |
| M3 | + (5) | ND | 8/15 | 2.14x10<sup>5</sup> (±1.05x10<sup>5</sup>) |
| M4 | + (9) | + | 5/15 | 2.3x10<sup>5</sup> (±1.05x10<sup>5</sup>) |
| M5 | + (4) | + | NA | NA |
| M6 | + (4) | + | NA | NA |
| M7 | + (4) | + | NA | NA |

DPF: days post tick feeding
ND: not determined because the animal was sacrificed for spirochete isolation
NA: not applicable because the ticks were kept alive for future experiments
<sup>‡</sup> as determined by qPCR

### Rates of filial infection in F1 nymphs

After ticks molted to the third instar stage, qPCR determined infection rates in 60 ticks (Table 1 and Figure S1). The remaining ticks were saved for future experiments. Out of the 60 F1 ticks, 23 were positive for *B. turicatae flaB* gene and filial infection rates were 38.3% (95%CI 26.3-51.8%). These findings determined vertical transmission rates in the offspring of ticks that autogenously reproduced.

## Discussion

This study arose from laboratory observations of *O. turicata* that were collected in a public park within a neighborhood of Austin, Texas. Ticks were reared from the nymphal stage to adulthood, and we observed autogenous reproduction. While autogeny has been reported in a close relative of *O. turicata, Ornithodoros parkeri* (8, 11), literature on the former is absent. The reasons for this lack of data are unclear. Without designed experiments housing individual ticks and tracking their molt progress, the phenomenon may have been overlooked during colony maintenance.

In Argasidae, nuances in autogeny depend on reproductive and feeding behaviors. Several species are fully autogenous, such as *Otobius* spp. and *Antricola* spp. In the adult stage these genera have vestigial mouthparts and never blood feed (12). *Alveonasus lahorensis* females are obligatory autogenous in their first gonotrophic cycle but have to secure a blood meal to produce subsequent egg batches (13). Facultative autogeny in the first gonotrophic cycle has been reported at variable rates in species currently classified in the genera *Ornithodoros, Argas*, and *Carios* (8, 10, 12, 14). Interestingly, in other arthropods like mosquitoes autogeny varies extensively between conspecific populations across distribution ranges (15). While studies suggest that genetic and environmental factors (e.g. temperature) modulate autogeny in soft ticks (11), additional work is needed in *O. turicata* to identify its frequency between geographically distinct populations.

We evaluated filial infection rates and successful *B. turicatae* transmission to mice from the autogenously produced cohort of *O. turicata*. These findings indicate that after the transovarial passage through ticks, *B. turicatae* remained infectious. Determining infectivity of *B. turicatae* to mice was important because prior work on continuous vertical transmission through successive generations of *Ornithodoros* species is ambiguous. For example, ∼22% of vertically infected cohorts of *Ornithodoros papillipes* failed to transmit *Borrelia sogdiana* in the eighth generation compared to ∼100% in earlier generations (16). Additionally, a complete loss of transmissibility of *Borrelia duttonii* by tick bite was reported by the fifth generation in transovarially-infected *Ornithodoros moubata* (17). However, Burdorfer and Varma failed to reproduce these findings using a different strain of spirochete and population of tick (18). With *B. turicatae*, spirochetes remained infectious to mice after vertical maintenance by the vector for five generations (7). In previous studies, the life stage where ticks became infected was known to the investigators. A caveat in our work was that *O. turicata* ticks were field collected and it was unclear if the infected parental tick acquired spirochetes vertically or from an infectious blood meal. We have isolated and cultured the *B. turicatae* strain from this study and will assess vertical transmission after infecting *O. turicata* at different life stages.

We determined spirochete densities in individual ticks to better understand transovarial transmission rates in the offspring of autogenously reproduced ticks. Our estimates of *B. turicatae* loads were based on *flaB* copies per tick, and initially appeared high. However, work with *Borrelia hermsii* and *Borrelia burgdorferi* indicate that the pathogens are polyploid having ∼16 chromosomal copies per cell depending on the bacteria’s growth stage (19-21). With this consideration, a more accurate estimation of *B. turicatae* densities in the ticks was likely ∼10-fold less, or ∼10^4^ spirochetes per tick.

While our study began to evaluate autogeny in *O. turicata* there were limitations. Our findings are for one strain of *B. turicatae*. It will be important to know the rates of vertical transmission between *B. turicatae* isolates given their genomic plasticity. RF spirochetes possess complex genomes with upwards of 15 plasmids, and isolates vary significantly in their plasmid content (4). Furthermore, in this study we evaluated one population of tick from Texas, while *O. turicata* possesses a wide distribution range. More elaborate studies will identify phenotypic differences between spirochete isolates and tick populations.

It is important to define the cost-benefit ratio of autogeny for both the tick vector and pathogen. Tick eggs and larvae are less resistant to adverse environmental conditions (13, 22); however, autogenous reproduction could maximize fitness in newly established tick populations with transient hosts. This is consistent with ecological bet-hedging (23). Indeed, facultative autogeny can accelerate population growth after dispersal into new suitable habitats in patchy landscapes, as has been proposed in triatomine kissing bugs (24). Additionally, newly colonized habitats containing *B. turicatae* infected larvae could facilitate a rapid expansion of enzootic foci. Studies by Davis highlighted the significance of *O. turicata* as a long-term reservoir of *B. turicatae* (7), and our findings support this notion. While more work is needed to delineate the mechanisms of vertical transmission, our work lays the foundation for studying *O. turicata* reproduction and spirochete-vector interactions.

## Materials and Methods

### Ethics statement

Animal studies were approved by the Baylor College of Medicine (BCM) Institutional Animal Care and Use Committee (protocol AN7086). All work and animal husbandry were in accordance to the United States Public Health Service policy on Humane Care and Use of Laboratory Animals and the Guide for the Care and Use of Laboratory Animals.

### Collection and speciation of ticks

In March 2022, field efforts were implemented to collect ticks. The ticks were baited with dry ice and housed at 24 ± 2 °C and 85% relative humidity (25). The species of tick was confirmed by microscopy and through PCR amplification of a fragment of the mitochondrial *cox1* gene using DNA from two ticks and the LCO and HCO primers (26).

The amplicons were purified using the Mag-Bind Total Pure NGS beads (Omega Bio-tek, Norcross, GA, USA) and sequenced. The sequences were deposited in GenBank.

### Tick feedings and detection of murine infection

To check for *Borrelia* infection, field-collected *O. turicata* were divided in pools of 8 to 10 ticks and fed on Institute for Cancer Research (ICR) mice, as previously described (4). Blood was collected for 10 consecutive days by tail nick and infection was determined by visualizing spirochetes using a dark field microscope (Zeiss Axio Imager A2, Oberkochen, Baden-Württemberg, Germany). After feeding, the ticks’ life stage was determined by visualization of the genital aperture. They were sorted into pairs of males and females, individual females, or groups of nymphs (genital aperture is not apparent at this developmental stage). Ticks were housed in 50 ml TubeSpin Bioreactor tubes (MidSci, St. Louis, MO, USA).

The F1 progeny from a single female was further evaluated. The remaining larvae from the other females that autogenously reproduced were kept for other studies. The ticks were reared by feeding on mouse pups and at the second instar stage they were tested for infection by feeding on seven ICR mice (15 to 18 ticks per animal; n=107). Infection was evaluated by microscopy, as stated above. Thirty days after tick feedings, all animals were euthanized and blood was collected to determine infection by evaluating seroconversion to *B. turicatae* protein lysates, as reported (4).

### Molecular detection of *B. turicatae* in F1 ticks

Filial infection rates were determined on a subset of progeny using a duplex qPCR assay targeting *B. turicatae flaB* and the *O. turicata B-actin* gene, as described (27). DNA was extracted from individual unfed third instar stage nymphs using the DNeasy Blood and Tissue kit (Qiagen, Hilden, Germany). Each sample was run twice in triplicate. Spirochete loads in each sample was determined as previously described (28). The Wilson interval with continuity correction was calculated for the resulting point estimate of filial infection rates to obtain 95% confidence interval using R (29).

## Acknowledgments

We would like to thank Alex Kneubehl and Bonny Mayes for their help in the collection of ticks. This work was supported by NIAID, NIH grants AI137412 and AI144187 (JEL).

## Figure Legends

**Figure S1**. Calculated *flaB* copies per tick. Twenty-three ticks were tested with duplex qPCR and box and whisker plots are shown with each data point representing an infected tick. Cohorts of ticks are shown according to the mouse (M1 -M4) they fed upon.

